# RNAi gene knockdown in the poultry red mite, *Dermanyssus gallinae* (De Geer 1778), a tool for functional genomics

**DOI:** 10.1101/2020.09.28.316380

**Authors:** Wan Chen, Kathryn Bartley, Francesca Nunn, Alan S. Bowman, Jeremy M. Sternberg, Stewart T. G. Burgess, Alasdair J. Nisbet, Daniel R. G. Price

**Affiliations:** Moredun Research Institute, Pentlands Science Park, Edinburgh EH26 0PZ, United Kingdom; Institute of Biological and Environmental Sciences, School of Biological Sciences, University of Aberdeen, Aberdeen AB24 3FX, United Kingdom

**Keywords:** RNA interference, poultry red mite, functional genomics, gene silencing, gene knockdown

## Abstract

**Background:** The avian haematophagous ectoparasite, *Dermanyssus gallinae* or the poultry red mite, causes significant economic losses to the egg laying industry worldwide and also represents a significant welfare threat. Current acaricide-based controls are unsustainable due to the mite’s ability to rapidly develop resistance, thus developing a novel sustainable means of control for *D. gallinae* is a priority. RNA interference (RNAi) mediated gene silencing is a valuable tool for studying gene function in non-model organisms, but is also emerging as a novel tool for parasite control.

**Methods:** Here we use an *in silico* approach to identify core RNAi pathway genes in the recently sequenced *D. gallinae* genome. In addition we utilise an *in vitro* feeding device to deliver dsRNA to *D. gallinae* targeting the *D. gallinae vATPase subunit A* (*Dg vATPase A*) gene and monitor gene knockdown using quantitive PCR (qPCR).

**Results:** We identified core components of the small interfering RNA (siRNA) and micro RNA (miRNA) pathways in *D. gallinae*, which indicate these gene silencing pathways are likely functional. Strikingly, the Piwi-interacting RNA (piRNA) pathway was absent in *D. gallinae*. In addition, we demonstrate that feeding *Dg vATPase A* dsRNA to adult female *D. gallinae* results in silencing of the targeted gene compared to control mites fed non-specific *lacZ* dsRNA. In *D. gallinae*, dsRNA mediated gene knockdown is rapid, detectable 24 hours after oral delivery of dsRNA and persisted for at least 120 hours.

**Conclusions:** This study has shown the presence of core RNAi machinery components in the *D. gallinae* genome. In addition, we have developed a robust RNAi methodology for targeting genes in *D. gallinae*, which will be of value for studying genes of unknown function and validating potential control targets in *D. gallinae*.

## Background

Poultry red mite (*Dermanyssus gallinae* (De Geer, 1778)), is an avian haematophagous ectoparasite with a worldwide distribution and a prevalence of 83% in European hen egg laying facilities (1). There are five life cycle stages in *D. gallinae*: egg, larvae, protonymph, deutonymph, and adult; and blood feeding is only a feature of the latter three stages (2, 3). This parasite lives off-host in the cracks and crevices in the hen facilities and only emerges to bite the host in darkness for a bloodmeal, which takes around 30 to 90 min during which each mite consumes ∼200 μg of blood per feed (3, 4). Thus, in severe infestations where each laying hen may be infested with up to 500,000 mites, infestation can lead to multiple behavioural and physiological changes in the birds, such as restlessness, irritation, anaemia, feather pecking, cannibalism, and increased mortality rates (5, 6). Also, *D. gallinae* has been reported as the vector for a number of bacterial and viral diseases of birds as well as zoonotic agents (7, 8). Apart from the hen welfare issues caused by *D. gallinae*, it also increases the operational expenditure for hen egg production through losses in feed conversion ratio, downgrading of eggs and decreased egg output (6, 9, 10). Overall, the estimated annual cost of *D. gallinae* (production loss plus costs of control) was estimated as €231 million in Europe in 2017 (6). Conventional control of *D. gallinae* is through the use of chemical acaricide treatments of poultry houses or through systemic acaricides administered via drinking water (11). However, with the increased incidence of resistance against some acaricides (12) and concerns over residues in food, multiple chemical treatments have been withdrawn from use in the EU (2).

Development of novel stratergies for control of *D. gallinae* is a priority and gene target identification for development of novel control approaches is facilitated by the recent publication of transcriptomes and the genome of *D. gallinae* (13, 14). One key tool for exploiting these genomic and transcriptomic resources for novel target identification is transcriptional silencing. Since the discovery of RNA interference (RNAi) as a tool for silencing gene expression in the free living nematode, *Caenorhabditis elegans* (15), its application has been widened into various fields including novel arthropod control strategies (16, 17). Irrespective of the organism in which transcriptional silencing is to be investigated, two essential components are required for successful RNAi: 1) the presence of a functional RNAi pathway and 2) an appropriate delivery method for the gene-specific double-stranded RNA (dsRNA) to initiate the silencing process. RNAi pathways are present in many of the mite species investigated to date (reviewed in (18)) and, for several species, delivery of dsRNA has been achieved through soaking mites in solutions containing the dsRNA (e.g. see (19, 20)). While the immersion method of dsRNA delivery has also caused gene knockdown in *D. gallinae* (21), high mortality rates were observed, thus hampering interpretation of transcriptional silencing data and necessitating the development of a better approach of dsRNA administration to *D. gallinae*.

Here, we use the recently published *D. gallinae* genome (13) and associated transcriptomic data to describe the RNAi pathway in *D. gallinae* and also investigate an optimised delivery method to ascertain the optimal properties of the dsRNA for RNAi in this species. RNAi mediated gene silencing was investigated in adult female *D. gallinae* mites by targetting *D. gallinae vacuolar ATPase subunit A (Dg vATPase A)*, which has previously been targeted in arthropods, including the two spotted spider mite, *Tetranychus urticae* (22).

## Methods

### Ethics approval

All procedures to gather samples from animals were approved by the Moredun Animal Welfare and Ethical Review Body (AWERB) and were conducted under the legislation of a UK Home Office License (reference PPL 60/03899) in accordance with the Animals (Scientific Procedures) Act of 1986.

### RNAi Pathway Gene Annotation

Core RNAi pathway components were identified in the *D. gallinae* genome by sequence similarity to RNAi pathway genes from *Drosophila melanogaster*. Twelve core RNAi pathway genes were selected from *D. melanogaster* that included Argonaute proteins, RNaseIII proteins and dsRNA binding proteins. Core *D. melanogaster* RNAi components were selected to represent three RNAi pathways, including: miRNA pathway (*argonaute1, Dicer 1, loquacious, Drosha, Pasha, Exportin-5*); siRNA pathway (*Argonaute2, Dicer 2, R2D2*) and piRNA pathway (*Aubergine, Piwi, Argonaute3*). Amino acid sequences for all twelve *D. melanogaster* RNAi pathway components were retreived from Flybase (23) and used as query for blastp searches against predicted peptides from the *D. gallinae* genome (13). Identified *D. gallinae* RNAi components were considered sequence orthologues for a given RNAi pathway component when they met the criteria of reciprocal best hit and included one-one and one-many sequence orthologues. Functional domains were identified in retrieved *D. gallinae* sequences using Pfam sequence searches (24).

### Phylogenetic analysis

For phylogenetic analysis selected argonaute-like protein sequences were aligned using MUSCLE (25). Short sequences (<50% of the protein’s consensus length) and predicted non-functional sequences due to absence of either PAZ domain (PF02170) or Piwi domain (PF02171) were removed from the alignment. All sequences used for phylogenetic reconstruction are shown in Additional file 1: Table S1. Ambiguously aligned positions were excluded by trimAL v1.2 (26) and a maximum-likelihood (ML) phylogenetic tree was constructed in MEGA 10.1.8 (27) using a LG+G substitution model. Statistical tree robustness was assessed using bootstrap analysis (1000 bootstrap replicates).

### Amplification of *D. gallinae vATPase A* gene for RNAi validation

The full-length sequence of *D. gallinae vacuolar ATPase subunit A* (*Dg vATPase A:* DEGAL4806g00010) was retrieved from the OrcAE database for *D. gallinae* (13, 28). To validate the OrcAE gene model, *Dg vATPase A* (DEGAL4806g00010) was used in a blastx search against NCBI nr protein database and sequences with high similarity were retrieved from: *Varroa destructor* (XP_022670783 and XP_022670784); *Tropilaelaps mercedesae* (OQR76956); *Galendromus occidentalis* (XP_003741079) and *Ixodes scapularis* (XP_029849202). All sequences were aligned using MUSCLE (25) and primers were designed based on conserved regions across *Dg vATPase A* and all other aligned sequences. The clustalx alignment and region used for primer design is shown in Additional file 2: Fig S1. The full-length coding sequence of *Dg vATPase A* was amplified using female *D. gallinae* cDNA as template and verified by Sanger sequencing.

### dsRNA Synthesis

Region 1 (R1: 495 bp, corresponding to exon 4 – 7) and Region 2 (R2: 385 bp, corresponding to exon 8) of the *Dg vATPase A* gene were amplified from cDNA generated from adult female *D. gallinae* using Phusion proof-reading polymerase (Thermo Fisher Scientific, Waltham, MA, USA). Each forward and reverse primer contained an NcoI and NheI restriction enzyme site, respectively, to allow directional cloning into the RNAi vector pL4440 (pL4440 was a gift from Andrew Fire [Addgene plasmid # 1654; http://n2t.net/addgene:1654 ; RRID:Addgene_1654]). Primer sequences are shown in Additional file 3: Table S2. Amplification products for *Dg vATPase A* R1 (495 bp) and R2 (385 bp) were digested with NcoI and NheI and cloned into the corresponding restriction enzyme sites of pL4440. Plasmids were used to transform chemically competent *E. coli* JM109 cells (Promega, Madison, WI, USA) and plasmid was isolated *E. coli* transformants using a Wizard® *Plus* SV Minipreps DNA Purification System (Promega, Madison, WI, USA). Both RNAi constructs containing *Dg vATPase A* R1 and R2 were verified by Sanger sequencing. For control (non-target) dsRNA production we used a previously generated construct containing a region of the *E. coli* strain K-12 *lacZ* gene NC_000913 (319bp; 63 – 381 bp of the CDS) cloned into SacI and SmaI sites of pL4440 (19). dsRNA was synthesized using the T7 RiboMAX Express RNAi System (Promega, Madison, WI, USA), according to the manufacturer’s instructions. For RNA synthesis *Dg vATPase A* pL4440 plasmids were linearized with either NcoI or NheI for sense or antisense transcription, respectively. Control *lacZ* pL4440 plasmid was linearized with SmaI or BglII for sense or antisense transcription, respectively. Purified dsRNA was quantified on a NanoDrop One spectrophotometer (Thermo Fisher Scientific, Waltham, MA, USA) and analysed by agarose/TAE gel electrophoresis to confirm quality and predicted size.

### siRNA Synthesis

*Dg vATPase A* siRNAs were synthesised by either *in vitro* digestion of long dsRNAs (Method-1) or chemical synthesis of 27-mer blunt dsRNAs (Method-2). Method-1: Long dsRNAs for R1 and R2 of the *Dg vATPase A* gene and *lacZ* control gene (120 μg of each dsRNA) were incubated with 0.2 units/μL ShortCut® RNase III (New England BioLabs, Ipswich, MA, USA) for 3 h at 37°C to produce a heterogeneous mix of short (18-25 bp) siRNA. Reactions were stopped with EDTA according to the manufacturer’s protocol, ethanol-precipitated and the size-distribution of digested RNAs validated by electrophoresis using a 4% agarose gel. Method-2: Dicer-substrate siRNAs (27-mer blunt dsRNAs) were designed based on the coding sequence of the *Dg vATPase A* gene and *lacZ* control gene using the Eurofins Genomics siMAX siRNA design tool and commercially synthesized and annealed by Eurofins Genomics (Eurofins Genomics, Ebersberg, Germany). The sequence of each siRNA is shown in Additional file 4: Fig S2.

### RNAi Feeding Trials

Mixed stage and mixed sex *D. gallinae* mites were collected from commercial egg-laying facilities and stored in vented 75 cm^2^ tissue culture flasks (Corning Inc, Corning, NY, USA) at room temperature for seven days, after which they were stored at 4°C for 3 weeks without access to food. For oral delivery of dsRNA and siRNA to *D. gallinae* mites, approx. 100 mites were housed in an *in vitro* feeding device constructed as described previously (29). Each replicate feeding device contained 200 µl of freshly-collected heparinised goose blood (20 units/ml) with dsRNA at concentrations described in each experiment. For each dsRNA feeding experiment, biological replicates consisted of independent group of mites in replicate feeding devices (n = 3 - 6 depending on experiment). Feeding devices were placed in a Sanyo MLR-351H relative humidity incubator for 3h at 39°C; 21h at 25°C both at 85% relative humidity. After 24 h, fed adult female mites were recovered from each replicate feeding divice and transferred to separate labelled 1.5ml tubes, which were held at 25°C in dark conditions for the duration of the experiment. Mites from each replicate group were flash-frozen in liquid nitrogen at time-points indicated in each experiment and stored at -70°C for later RNA extraction.

### Quantitative real-time PCR (qPCR) analysis

Real-time quantitative polymerase chain reaction (qPCR) was used to quantify *Dg vATPase A* gene expression in adult female mites from RNAi feeding trials. Mites were homogenized with a tube pestle and total RNA was isolated using an RNeasy® plus micro kit (Qiagen, Hilden, Germany) which included a gDNA eliminator spin-column. Total RNA was quantified using a NanoDrop One (Thermo Fisher Scientific, Waltham, MA, USA) and first-strand cDNA synthesized using a QuantiTect® Reverse Transcription Kit (Qiagen, Hilden, Germany), according to the manufacturer’s protocol.

qPCR primers were designed using Primer3Plus (30), primer sequences are shown in Additional file 3: Table S2. Primers were checked for specificity by alignment of *D. gallinae* target sequences with goose *Anser cygnoides v-ATPase A* (XM_013196364) and *GAPDH* (XM_013199522). In addition, qPCR products were sequenced demonstrating amplification of only *Dg vATPase A* and *D. gallinae GAPDH*. For construction of standard curves, qPCR primers were used to amplify *Dg vATPase A* (DEGAL4806g00010) and *GAPDH* (DEGAL4146g00090) from adult female *D. gallinae* cDNA. Amplification products were cloned into pJET1.2 (Thermo Fisher Scientific, Waltham, MA, USA) and verified by DNA sequencing. Plasmids were used in qPCR experiments to construct standard curves from 10^2^–10^8^ copies of each gene. Ten microlitre qPCR reactions comprised 1× PowerUp SYBR green master mix (Thermo Fisher Scientific, Waltham, MA, USA), 500 nM of forward and reverse primers, and cDNA derived from 1 ng total RNA for each sample. PCR reactions were performed on an Applied Biosystems 7500 Real Time PCR System; thermal cycling conditions were 50°C for 2 min, 95°C for 2 min, followed by 40 cycles at 95°C for 15 s, 58°C for 15 s, and 72 °C for 1 min. *Dg vATPase A* gene expression was normalized to housekeeping gene *GAPDH* and expression levels reported relative to control (*lacZ*) dsRNA fed mites. qPCR experiments were performed in triplicate and included no template controls and no reverse transcription controls with each run.

### Statistical analyses

Analysis of *Dg vATPase A* gene expression levels in RNAi feeding trials were performed using GraphPad Prism version 8.0.0 for Windows (GraphPad Software, La Jolla, CA, USA). Datasets were analysed using either Student’s *t-test* or one-way ANOVA with Dunnett’s multiple comparison test (as indicated) and p-values of <0.05 were considered significant.

## Results

### miRNA and siRNA pathways are present in the *D. gallinae* genome

We used a systematic search for core RNA interference (RNAi) genes involved in the siRNA pathway (*Argonaute2, Dicer 2, R2D2*); miRNA pathway (*argonaute1, Dicer 1, loquacious, Drosha, Pasha, Exportin-5*); and piRNA pathway (*Aubergine, Piwi, Argonaute3*) in *D. gallinae*. Our searches identified *D. gallinae* orthologues of at least one core gene in the siRNA and miRNA and pathways (See both Fig 1 and Table 1) and did not identify piRNA pathway orthologues, suggesting that this pathway is not present in *D. gallinae*.

**Table 1.**
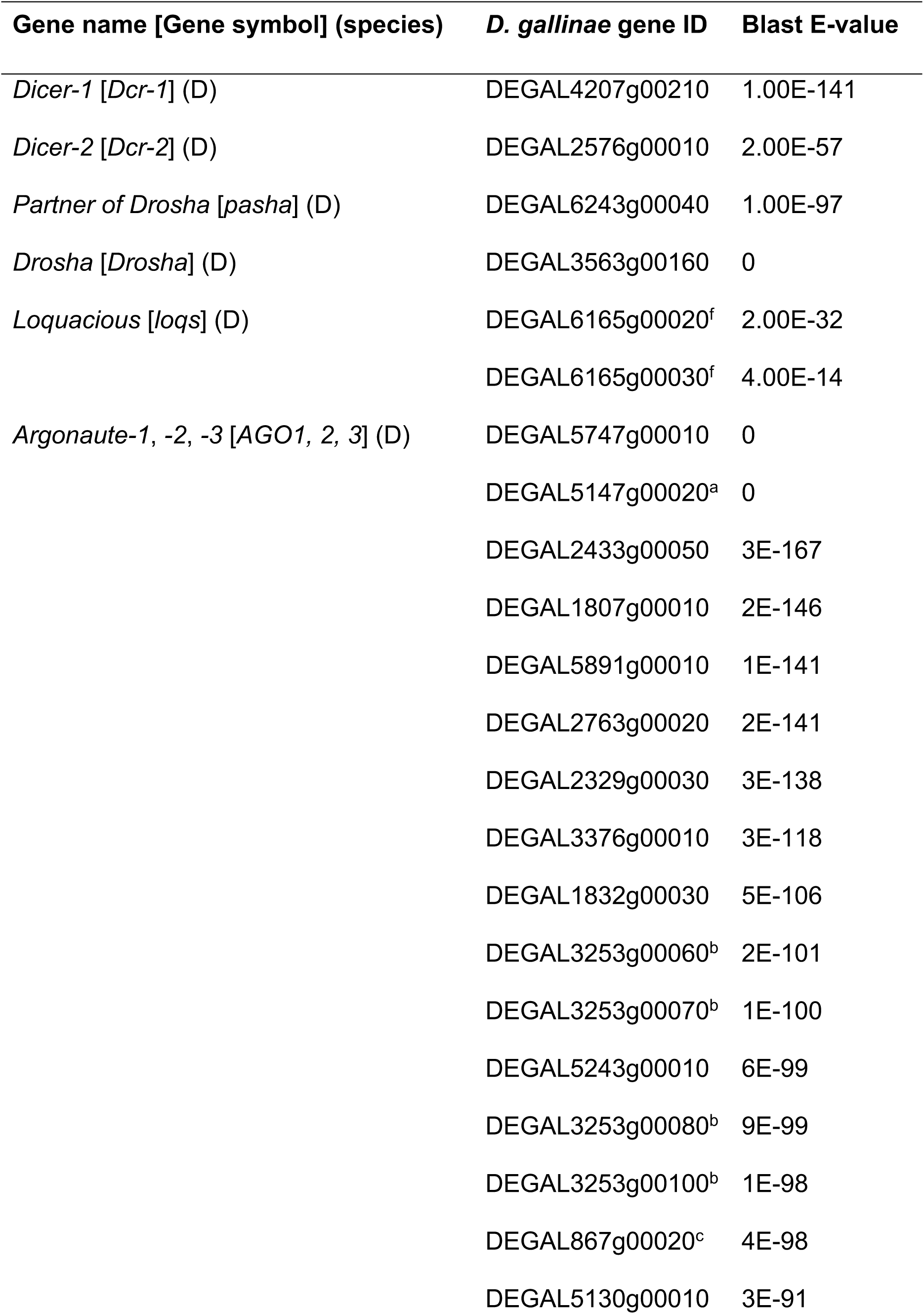
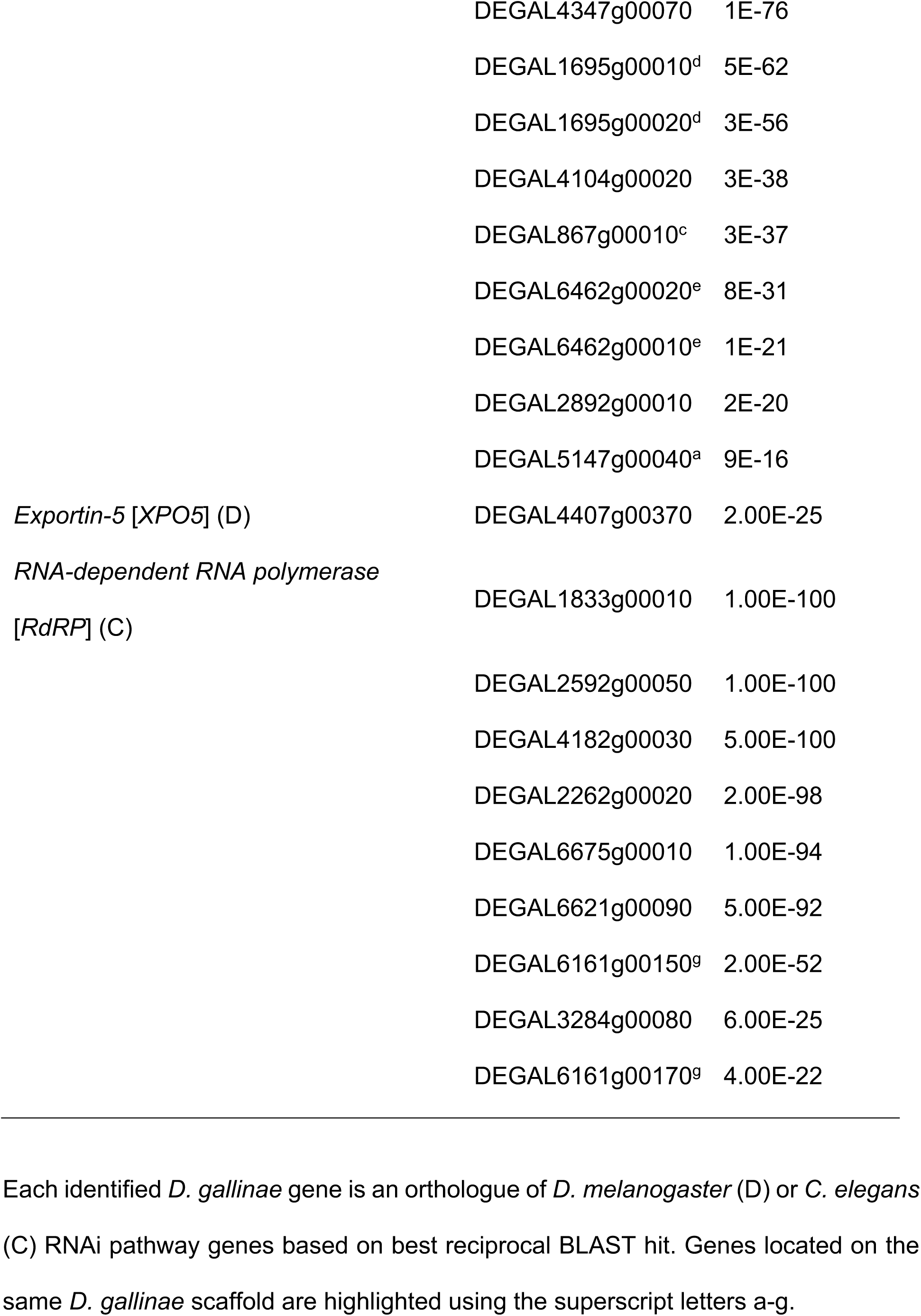
Identification of *D. gallinae* core RNAi pathway genes.

**Fig. 1.**
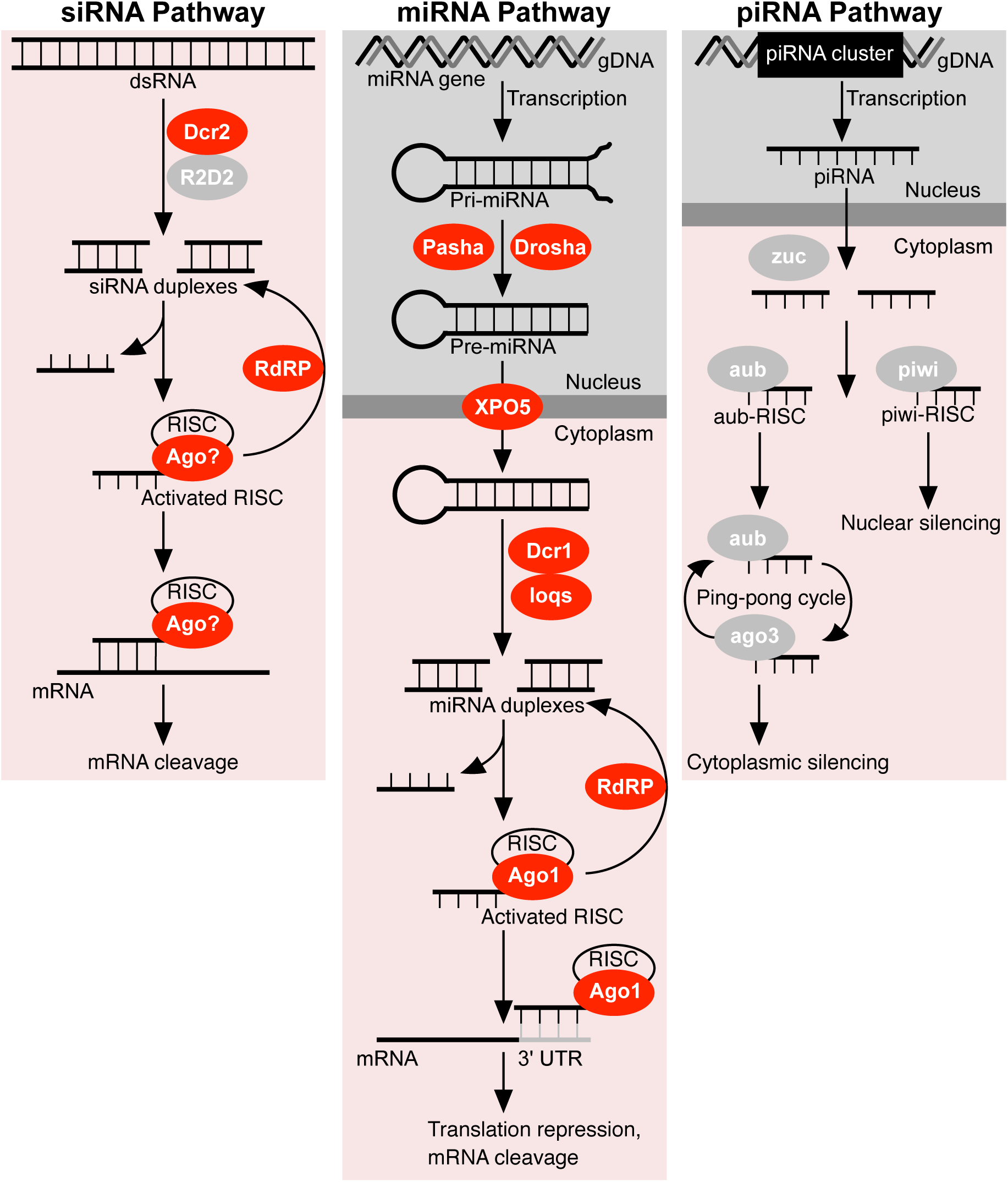
RNA interference (RNAi) pathways in *D. gallinae*. Core RNAi pathway enzymes are shown in either red (present in *D*.*gallinae*) or grey (absent in *D. gallinae*). **siRNA pathway:** dsRNA (either viral or experimentally introduced) is processed by Dicer-2 (Dcr-2) into 21-23 nt siRNAs and loaded into the RISC complex. Argonaute (Ago) cleaves the passenger strand of the siRNA and retains the guide strand which guides the active siRISC complex to the target mRNA. Full complementarity between the guide siRNA strand and target mRNA leads to cleavage of the mRNA. **miRNA pathway:** The miRNA gene is transcribed in the nucleus to generate pri-miRNA which is then cleaved by Drosha and Pasha to form a pre-miRNA. The pre-miRNA is transported to the cytoplasm through Exportin-5 (XPO5) and cleaved by Dicer-1 (Dcr-1) to yield miRNA. The miRNA is loaded into the RISC complex and argonaute (Ago) cleaves the passenger strand of the miRNA and retains the guide strand which guides the active miRISC complex to the target mRNA. Partial complementarity between the guide strand and target mRNA leads to either translation repression or cleavage of the mRNA. miRNAs usually target several genes, with shared sequences in the 3’ untranslated region (UTR). In both pathways siRNAs and miRNAs are amplified by RNA-dependent RNA polyrmerase (RdRP). **piRNA pathway:** The piwi-interacting RNA (piRNA) pathway functions in germline cells to protect against transposable elements. Antisense piRNAs are transcribed from repetitive elements in gDNA and processed by zucchini (zuc) into 26 – 32 nt primary piRNAs. Primary piRNAs associate with either piwi or abuergine (Aub). Piwi associated piRNAs are translocated to the nucleus, while Aub associated piRNAs cleave cytoplasmic transposon transcripts and trigger a ‘ping-pong’ piRNA amplification. Following transposon transcript cleavage argonaute 3 (AGO3) is loaded with secondary piRNAs which in turn produce piRNAs that associate with Aub, resulting in silencing of cytoplasmic transposon transcripts.

### siRNA Pathway genes

Of the three currently known core siRNA pathway genes from *Drosophila* (*Argonaute2, Dicer2* and *R2D2*), we identified an orthologue of *Dicer2* in *D. gallinae* (DEGAL2576g00010; [Table 1]) which is required to cleave and yield a mature siRNA. Domain analysis of *D. gallinae* Dcr-2 revealed a similar domain architecture to the well-characterised *Drosophila* Dcr-2, although the PAZ domain was absent in *D. gallinae* Dcr-2 (Additional file 5: Fig S3). Gene expression data available through OrcAE shows that *D. gallinae Dcr-2* is universally expressed in adult male and female mites and all other life-stages. Our searches did not identify any other core siRNA pathway genes in *D. gallinae* and, while our searches identified a large Ago family (containing 25 members), it did not include a *D. gallinae Ago2* orthologue.

### miRNA Pathway genes

We identified orthologues of all six core miRNA pathway genes in *D. gallinae*. Identified *D. gallinae* miRNA pathway genes included: *<u>Drosha</u>* (DEGAL3563g00160; [Table 1]) and *<u>Pasha</u>* (DEGAL6243g00040; [Table 1]) required for miRNA biosynthesis. *<u>Exportin-5</u>* (DEGAL4407g00370; [Table 1]) required for export of pre-miRNA from the nucleus to cytoplasm. *<u>Dicer-1</u>* (DEGAL4207g00210; [Table 1]) and two copies of its binding partner *<u>Loquacious</u>* (DEGAL6165g00020; DEGAL6165g00030; [Table 1]) required to cleave and yield a mature miRNA. *<u>Argonaute-1</u>* (DEGAL5747g00010; DEGAL5147g00020; [Table 1]) required to target and slice complementary RNA transcripts.

Domain analysis of *D. gallinae* Dcr-1 (Additional file 5: Fig S3) and *Argonaute-1* ortholouges (Additional file 6: Fig S4.) demonstrate the presence of functional domains associated with activity. In addition, gene expression data available through OrcAE shows that all identified *D. gallinae* miRNA components are universally expressed in adult male and female mites and all other life-stages.

### Piwi and Ago2 proteins are absent in the *D. gallinae* genome

*D. gallinae* Ago/Piwi coding sequences were identified based on similarity to two Ago proteins (Ago1 and Ago2) and three Piwi proteins (aub, piwi, Ago3) from *Drosophila*. All Ago/Piwi sequence orthologues identified in *D. gallinae* met the criteria of reciprocal best-hit in *D. gallinae* and *Drosophila* genomes. Using this search methodology we identified 25 Ago orthologues in the *D. gallinae* genome, while no Piwi orthologues were identified (Table 1). Additional searches were performed, using the same search criteria, with Ago/Piwi protein coding sequences from *T. urticae*, a related mite species with a high-quality annotated genome (31). These additional searches did not identify any additional Ago orthologues in *D. gallinae*, and again failed to identify any Piwi coding sequences.

All functional Ago proteins contain two common structural features: PAZ domain (responsible for small RNA binding) and Piwi domain (responsible for catalytic activities) (32). Therefore, in order to identify potential functional *D. gallinae* Ago proteins, all 25 *D. gallinae* Ago orthologues were analysed for domains using the Pfam database (24). These searches identified 16 *D. gallinae* Ago orthologues that contained both PAZ domain (Pfam, PF02170) and Piwi domain (Pfam, PF02171) (Additional file 6: Fig S4). In addition, all 16 *D. gallinae* Ago orthologues have a DEDH catalytic slicer motif within each Piwi domain indicating that these Agos likely retain slicer activity (33).

*D. gallinae* Ago orthologues with both PAZ and Piwi domains (a total of 16 *D. gallinae* Agos) were compared with orthologues from other arthropods including: the two-spotted spider mite, *T. urticae* (6 Ago orthologues and 7 Piwi orthologues) and insects (as detailed in Fig. 2). Our phylogenetic analysis identified two *D. gallinae* Ago1 orthologues (DEGAL5147g00020; DEGAL5747g00010) likely to be involved in the miRNA pathway. The remaining *D. gallinae* Ago orthologues belong to two major clades, the first containing seven members and are closely related to Ago1 proteins (Fig. 2, Ago1-like clade). The second major clade contains seven members and is unique to *D. gallinae* (Fig. 2, Dg Ago clade). Members of the Dg Ago clade show evidence of duplication, with four members (DEGAL3253g00060; DEGAL3253g00070; DEGAL3253g00080; DEGAL3253g00100) present as a tandem array on the same genomic scaffold. Strikingly, none of the identified *D. gallinae* Agos belong to either the Ago2 clade or Piwi clade (Fig. 2).

**Fig. 2.**
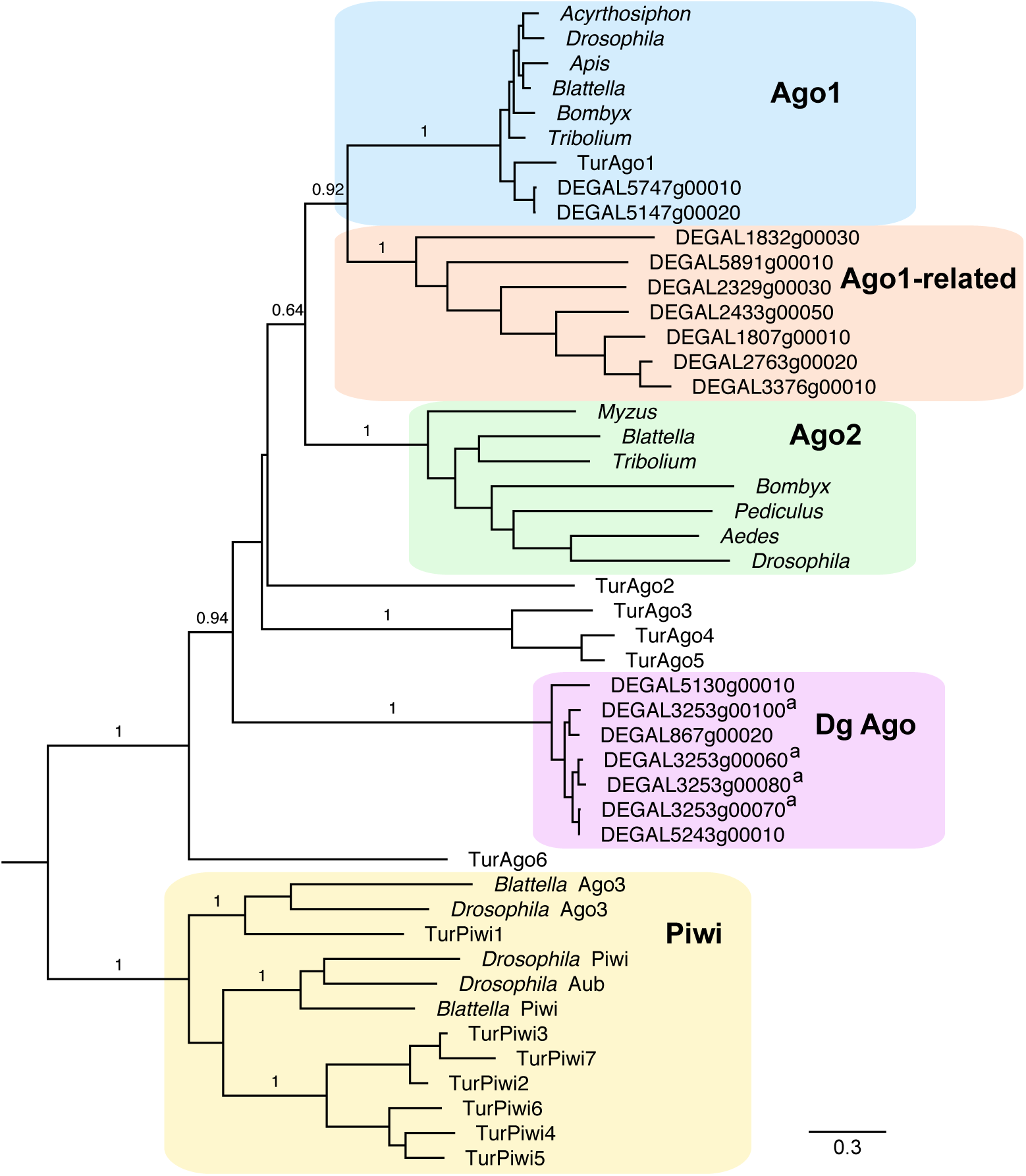
Phylogenetic analysis of Argonaute-like proteins from *D. gallinae* and other arthropods. All *D. gallinae* sequences are available at OrcAE using DEGAL accession numbers shown in the alignment. The species and accession number of all other sequences are shown in Additional file 1: Table S1. All protein sequences were aligned using MUSCLE and phylogenies reconstructed using maximum-likelihood (ML) methods with a LG+G substitution model. Bootstrap support values >0.6 from 1000 replicates are shown at each major node. Scale bar represents 0.3 substitutions per amino acid site. *D. gallinae* Argonaute genes that are located on the same genomic scaffold are indicated by a superscript letter.

### RNA-dependent RNA Polymerase (RdRP) is present in *D. gallinae* genome

RNAi is a conserved gene silencing mechanism in eukaryotes. In non-animal eukaryotes RNAi mediated gene silencing requires RNA-dependent RNA polymerase (RdRP) proteins (34). However, in the animals investigated thus far, only *C. elegans* requires RdRP for RNAi mediated gene silencing (34). Using *C. elegans* RdRP protein sequence as a query in blastp search against *D. gallinae* we identified 9 RdRP orthologues in the *D. gallinae* genome, that meet the criteria of reciprocal best hit in *D. gallinae* and *C. elegans* genomes (Table 1). Based on gene expression data available through OrcAE, all 9 *D. gallinae* RdRP orthologues are expressed and therefore may play a role in amplification and propagation of silencing signals (Table 1).

### Functional RNAi gene silencing in *D. gallinae*

Two regions of the *Dg vATPase A* gene were selected for synthesis of dsRNA. Region 1 (R1: 495 bp, corresponding to exon 4 – 7) and Region 2 (R2: 385 bp, corresponding to exon 8) were used for *in vitro* synthesis of dsRNA (Fig. 3). dsRNAs were incorporated into goose blood, which was delivered to adult female *D. gallinae* mites using an *in vitro* feeding device. After feeding, engorged mites were selected based on a visible blood meal contained within the abdomen and were used for gene expression analysis. Mites with no visible signs of feeding were discarded and not used for expression analyses. Feeding R1 and R2 dsRNA (both at 100 ng/μl) in separate feeding trials to adult female *D. galline* mites resulted in a significant 1.9-fold reduction (for both R1 and R2 feeding trials) in expression of *Dg vATPase A* compared with control mites that fed on blood containing non-specific *lacZ* dsRNA [*P*<0.05, one-way ANOVA with Dunnett’s multiple comparison test] (Fig. 4a). Under similar feeding methodology, combining R1 and R2 *Dg vATPase A* (100 ng/μl dsRNA, consisting of an equimolar mix of R1 and R2 dsRNA) dsRNAs resulted in a significant 2.6-fold reduction in expression of *Dg vATPase A* compared with control mites that fed from non-specific *lacZ* dsRNA [*P*<0.05, Student’s t-test] (Fig. 4b). During both feeding trials there was no gross observable phenotypic difference between mites fed with *Dg vATPase A* dsRNA and mites fed with control *lacZ* dsRNA.

**Fig. 3.**
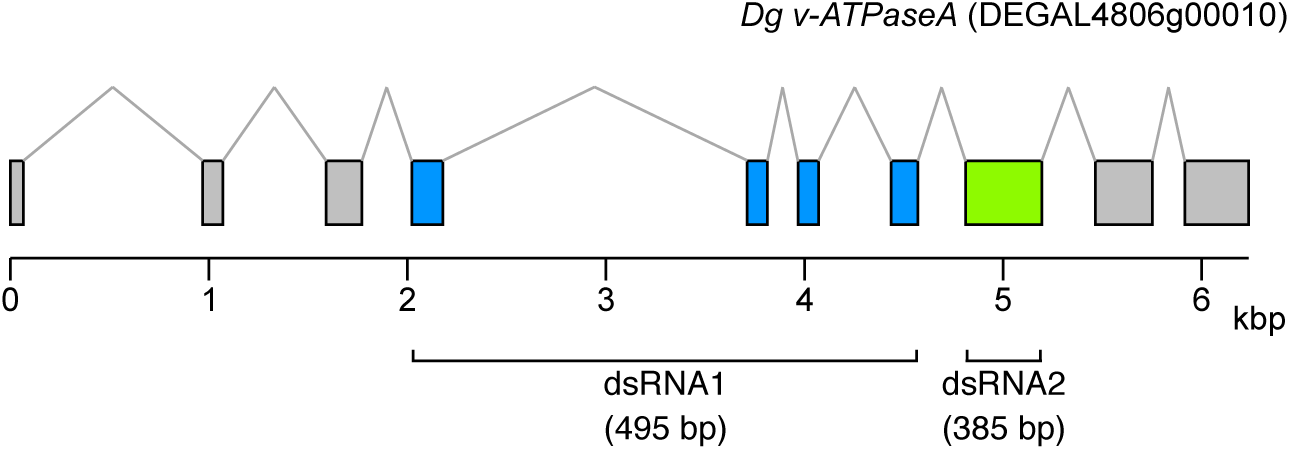
Regions of *Dg vATPase A* gene used for dsRNA synthesis. Schematic representation of the *Dg vATPase A* gDNA locus in *D. gallinae* gDNA scaffold DEGAL4806g00010 (34.1 kbp) location of Region 1 (R1) [exons 4, 5, 6 and 7] and Region 2 (R2) [exon 8] used for dsRNA synthesis.

**Fig. 4.**
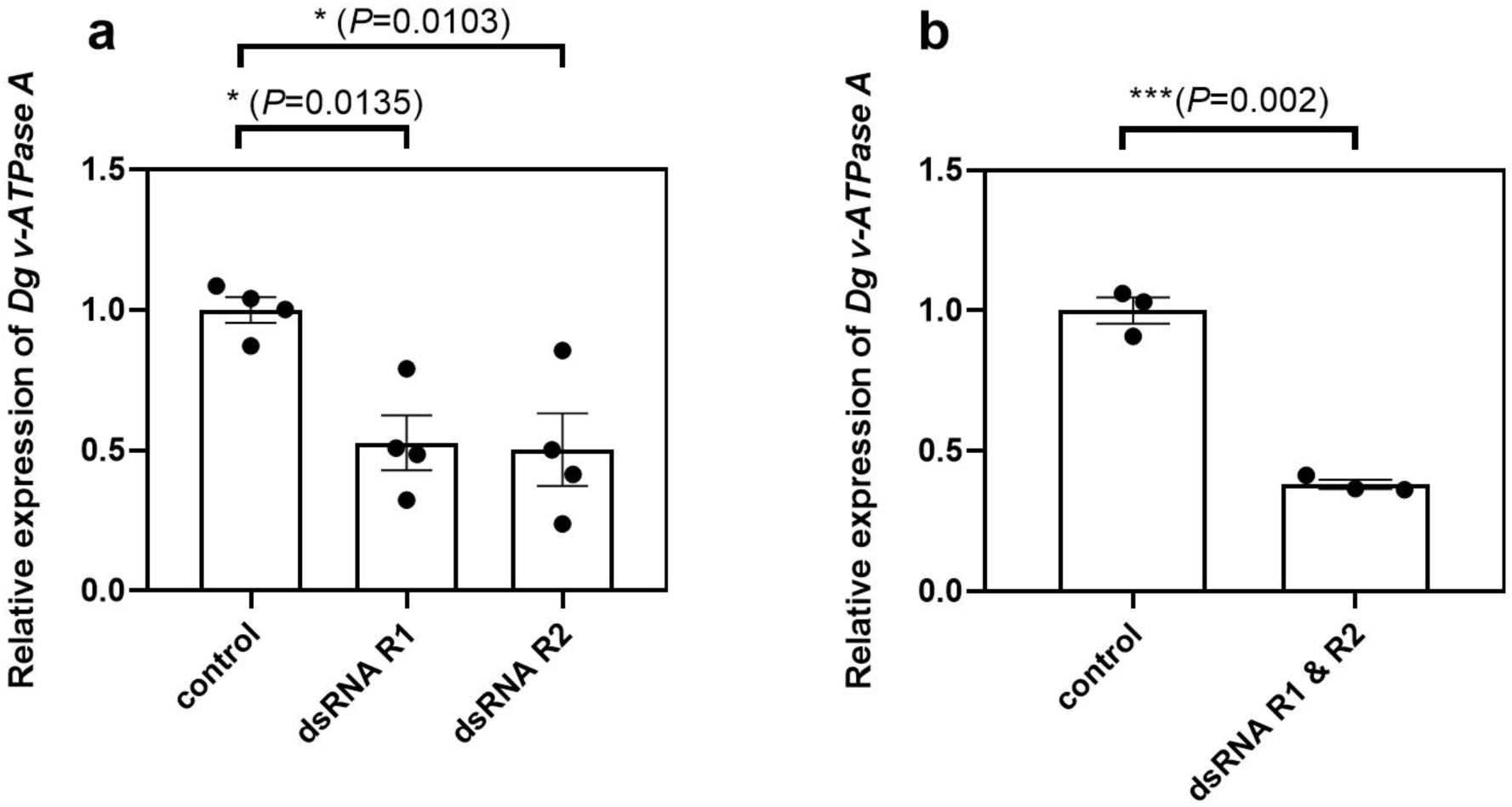
RNAi gene knockdown of *Dg vATPase A* in *D. gallinae*. qPCR gene expression analysis of *Dg vATPase A* expression in adult female *D. gallinae* mites fed on *lacZ* dsRNA (control) or *Dg vATPase A* dsRNA from Region 1 (R1) or Region 2 (R2). R1 and R2 *Dg vATPase A* dsRNAs were either fed separately **a** or combined **b** with a total dsRNA at 100 ng/μl (final concentration, with experiment **b** containing equimolar amounts of R1 and R2 dsRNA) in each experiment. *Dg vATPase A* expression shown at 96h post-feed and is normalised to *GAPDH* expression. Individual data points for biological replicates are shown with mean ± SEM indicated (n=3-4). Asterisks represent significant difference (*P*<0.05) between treatments determined by **a** one-way ANOVA with Dunnett’s multiple comparison test or **b** Student’s t-test.

### RNAi mediated gene silencing in *D. gallinae* is initiated quickly and is long lasting

Feeding combined R1 and R2 *Dg vATPase A* dsRNA to adult female *D. gallinae* mites resulted in rapid reduction of *Dg vATPase A* gene expression compared to control mites treated with the non-specific *lacZ* dsRNA. A significant reduction in *Dg vATPase A* expression was detectable by 24h post dsRNA delivery, and was maintained for at least 120h post dsRNA delivery (Fig. 5). At each time-point analysed, including 24h, 48h, 72h and 120h expression of *Dg vATPase A* was significantly reduced by 2-fold, 3.7-fold, 2.9-fold and 3.4-fold, respectively, relative to control (lacZ) fed mites [*P*<0.05, one-way ANOVA with Dunnett’s multiple comparison test] (Fig. 5).

**Fig. 5.**
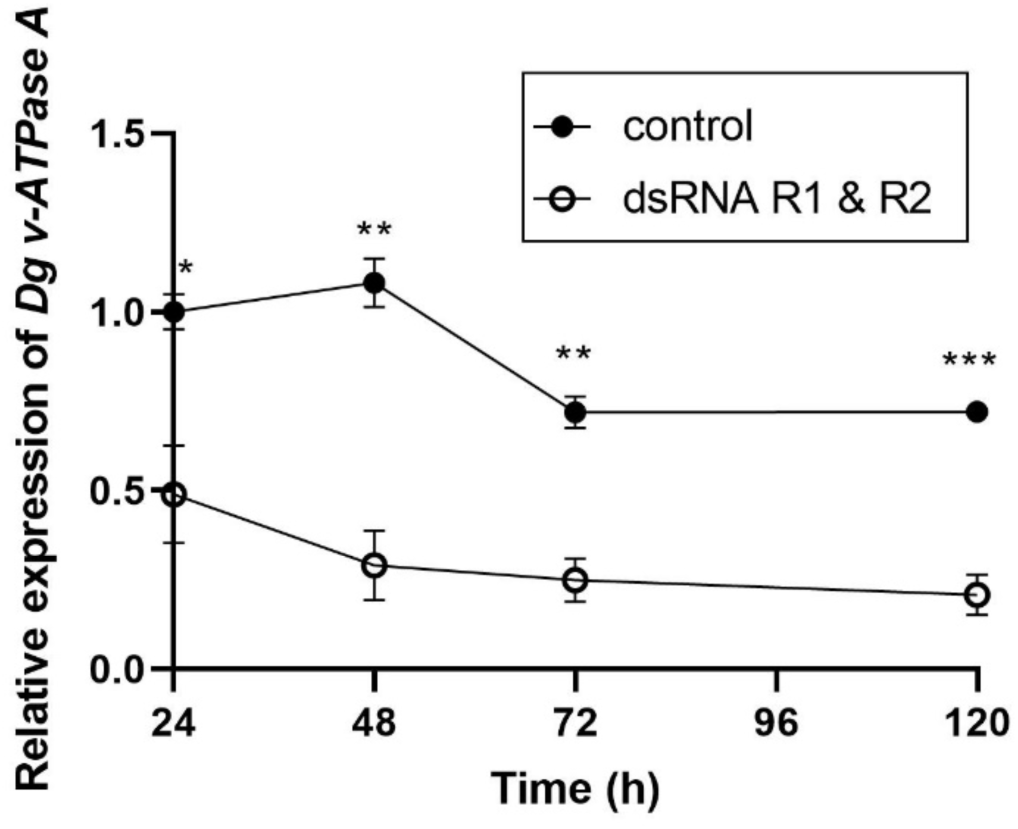
Persistence of RNAi gene knockdown of *Dg vATPase A* in *D. gallinae*. qPCR gene expression analysis of *Dg vATPase A* expression in adult female *D. gallinae* mites fed on *lacZ* dsRNA (control) or combined *Dg vATPase A* dsRNA from Regions 1 and 2 (R1 and R2). In control and *Dg vATPase A* dsRNA feeding experiments total dsRNA was delivered at 100 ng/μl (final concentration, containing equimolar amounts of R1 and R2 dsRNA) and *Dg vATPase A* expression levels monitored over a 120h post-feed time course. Individual data points for biological replicates are shown with mean ± SEM indicated (n=3-4). Asterisks represent significant difference (*P*<0.05) between treatments determined by one-way ANOVA with Dunnett’s multiple comparison test

### Oral delivery of siRNAs does not down-regulate target gene expression

Short dsRNAs were produced by either *in vitro* dicer treatment of synthesized long R1 and R2 dsRNAs (Method-1) or commercial synthesis of two 27 bp dsRNA corresponding to regions of the *Dg vATPase A* gene (Method-2). The sequence of each synthesized siRNA is shown in Additional file 4: Fig S2. Feeding trials were used to deliver either diced-R1/R2 dsRNAs (Fig. 6a) or synthesized 27 bp dsRNAs (Fig. 6b), along with the appropriate control dsRNA to adult female *D. gallinae* mites. Neither diced R1/R2 dsRNAs nor synthesized 27 bp dsRNAs resulted in down regulation of the *Dg vATPase A* gene expression, and treated mites had comparable expression levels with control fed mites (*P*=0.1073 for diced R1/R2 dsRNAs; *P*=0.6225 for synthesized 27 bp dsRNAs trial; Student’s t-test).

**Fig. 6.**
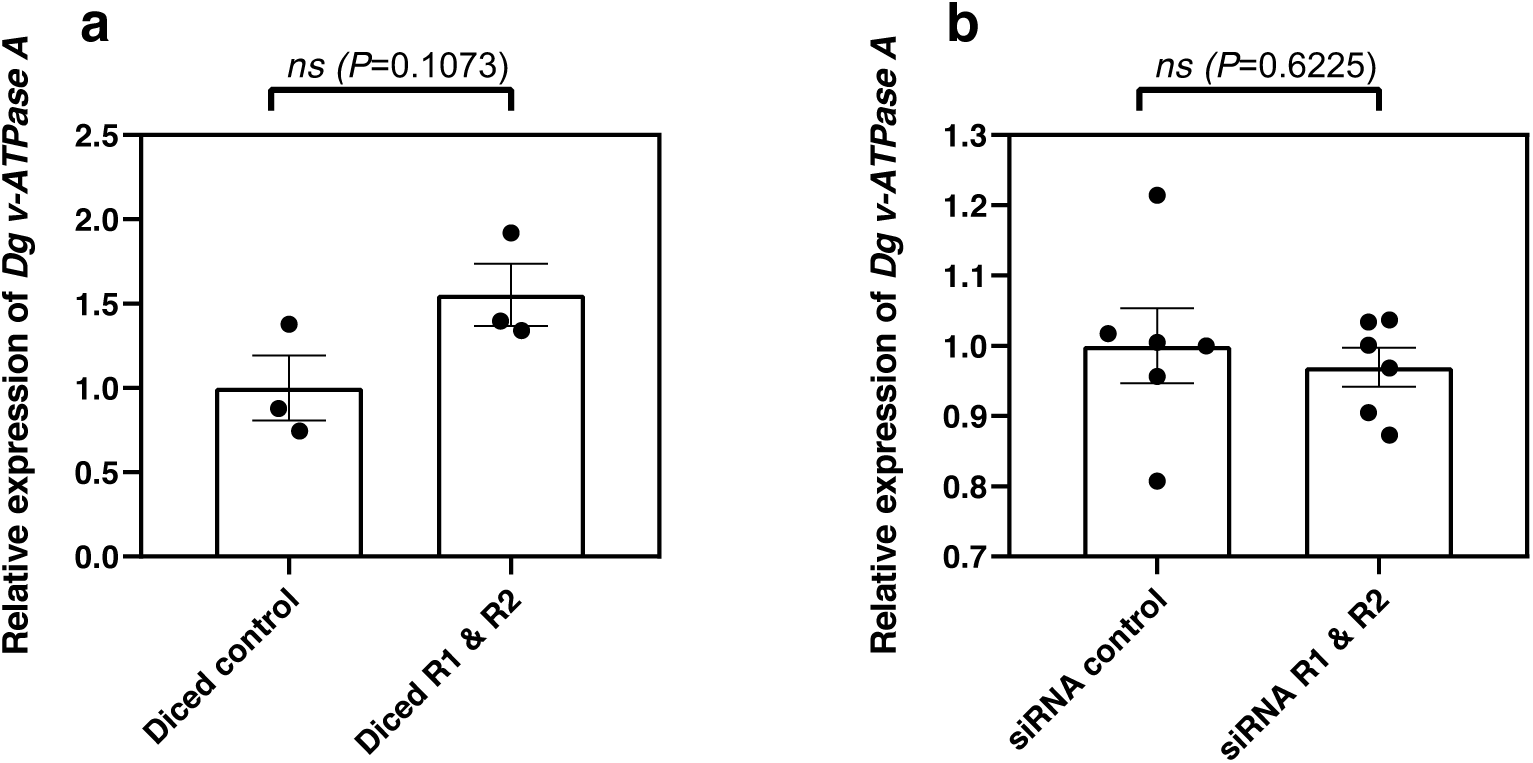
siRNAi gene knockdown of *Dg vATPase A* in *D. gallinae*. qPCR gene expression analysis of *Dg vATPase A* expression in adult female *D. gallinae* mites fed on *Dg vATPase A* siRNAs or *lacZ* siRNA (controul. **a** Long R1 and R2 *Dg vATPase A* dsRNAs and long *lacZ* dsRNA were for diced to produce short siRNAs and fed at 100 ng/μl. **b** siRNAs (27-mer) for *Dg vATPaseA* and *LacZ* (control) were commercially synthesized and fed at 100 ng/μl. For both **a** and **b** *Dg vATPase A* expression shown at 96h post-feed and is normalised to *GAPDH* expression. Individual data points for biological replicates are shown with mean ± SEM indicated (n=3-6). No significant differences (ns) were detected between treatment groups using Student’s t-test.

## Discussion

Here, we demonstrate the presence of genes encoding components of two of the main RNAi pathways in the genome of *D. gallinae*, namely the siRNA and miRNA pathways. Both exogenous and endogenous siRNA pathway components were identified in the *D. gallinae* genome but the major piRNA components, *Piwi, Aub* and *Ago3* genes, were lacking indicating either very dissimilar sequences of these genes or, more probable, the absence of this pathway in *D. gallinae*. We also demonstrate the utility of an *in vitro* feeding methodology (29) for the successful delivery of dsRNA to initiate prolonged, gene-specific RNAi in *D. gallinae* but show that orally-delivered short siRNA failed to initiate RNAi for the gene investigated herein.

In arthropods, three main RNAi mechanisms co-exist (small interfering RNAs (siRNA), microRNAs (miRNA), and Piwi-interacting RNAs (piRNA) (17) but in several species of the Acari investigated to date, the piRNA pathway has been notably absent. The piRNA pathway appears to be absent in: the sheep scab mite, *Psoroptes ovis* (19), the human itch mite, *Sarcoptes scabiei* (18) and the house dust mites, *Dermatophagoides farinae* (35), *Dermatophagoides pteronyssinus* (18) and *Euroglyphus maynei* (18). In *Drosophila*, where the piRNAi biology is best characterised, the piRNA pathway silences transposons during germline development, thereby protecting the inherited genome from mutation (36). The apparent lack of the piRNA pathway in *D. gallinae* and other members of the Acari, reflects the dynamic nature of RNAi pathways in mites and may indicate species-specific biology. The molecular mechanism/s which protect *D. gallinae* and other Acari lacking the piRNA pathway against the deleterious effects of transposon activity in the germline await investigation.

Our work presented here demonstrated that components of the siRNA pathway are present in the *D. gallinae* genome. One notable absence was the lack of a *D. gallinae* Ago2 orthologue, onto which the mature siRNA is loaded and is required for gene silencing (Fig 1). Although a definitive orthologue of Ago2 was not discovered in *D. gallinae*, the argonaute family was expanded with 25 family members, some of which were unique to *D. gallinae*, raising the possibility that, in *D. gallinae, Ago2* is replaced by another yet uncharacterised argonaute. Our experimental work presented here confirms that although an Ago2 orthologue is missing in *D. gallinae*, the siRNA pathway is functional and can be exploited be feeding exogenous dsRNA, resulting in specific knockdown of the targeted *Dg vATPase A* gene. In our RNAi feeding experiments, *Dg vATPase A* was chosen as a target, as it has been previously targetted in *T. urticae*, where down-regulation resulted in a colour change phenotype in *vATPaseA* gene silenced *T. urticae* mites (22). Using a similar dsRNA feeding methodology, we achieved gene silencing of *Dg vATPase A*, but this was not associated with any notable phenotypic changes. In experiments investigating dsRNA length, we were able to achieve robust and reproducible *Dg vATPase A* knockdown using dsRNAs of 385bp and 495bp. However, when we targeted the same gene with short siRNAs, produced from either dicing the long dsRNAs or commercially synthesizing a short 27bp siRNA, the gene knockdown effect was lost. Recent RNAi experiments in *T. urticae* demonstrate there is a size threshold for effective gene silencing, where long dsRNAs resulted in robust gene silencing, while shorter dsRNAs (100 – 200 bp) were not effective (37). In *D. gallinae* the lower limit of dsRNA size for efficient gene silencing is currently unknown and awaits further investigation. The lack of gene silencing by short siRNAs in *D. gallinae* may result from three possibilities: i) uptake of short dsRNA from food is limited and therefore does not induce RNAi pathway; ii) the abundance of siRNAs saturates the intracellular RNA uptake system system; or iii) short dsRNAs are transported but not recognized by cellular RNAi processing machinery.

Functional RNAi is an important tool in both model and non-model organisms to investigate genes of unknown function. For example, RNAi has been exploited in *Drosophila* to investigate gene function in cultured cells using high throughput screening methods (38). Functional RNAi in *D. gallinae*, using methodologies described here, is particularly timely for investigating genes of unknown function in *D. gallinae*. Recent completion of the draft *D. gallinae* genome sequence (959 Mbp assembly) identified 14,608 protein coding genes, of which 768 appear to be unique to *D. gallinae*, without similarity to proteins in the NCBI nr protein database (13). Therefore development of a robust and repoducible RNAi methodology, coupled with -omic technologies offers a powerful approach to begin investigating gene function and biology that is unique to *D. gallinae*. In addition, RNAi methodologies for *D. gallinae* are likely to be a useful tool in the development of novel control methods, including vaccine development, as it will support the research community with interests in *D. gallinae* control (39-41). RNAi mediated gene silencing in *D. gallinae*, coupled with either *in vitro* bioassays (29), or on-bird feeding assays (42) will allow rapid screening of potential *D. gallinae* vaccine candidates and druggable targets. Significantly, utilising RNAi to screen for vaccine candidates in *D. gallinae* will likely speed up the process of antigen discovery and in turn conform to 3R principles of using fewer animals for vaccine trials.

## Conclusions

We found evidence for the presence of two RNAi pathways in *D. gallinae* and also successfully demonstrated that our functional gene knockdown protocol can be initiated by feeding gene-specific dsRNA in an improved *in vitro* feeding device. This opens the door for larger scale, genome-wide screening for novel *D. gallinae* control targets and also the opportunity to ascribe functions, using phenotypic assays, to the multiple genes of unknown function identified within the *D. gallinae* genome, for which no homologues exist in other species.

## Availability of data and material

The *Dg vATPase A* nucleotide coding sequence is available in the NCBI database using the following accession number: MW032475.

## Acknowledgements

The authors gratefully acknowledge funding for this project from the Scottish Government Rural Affairs, Food and the Environment (RAFE) Strategic Research Portfolio 2016-2021. DRGP is supported by a research fellowship provided by the Moredun Foundation. WC is supported by a studentship provided by the Univeristy of Aberdeen and the Moredun Foundation. We would like to thank the Bioservices Unit, Moredun Research Institute, for expert care of the animals.

## Authors’ contributions

AJN, DRGP, STGB, WC designed the study; DRGP, WC performed research; AJN, DRGP, STGB, WC drafted the paper. ASB, FN, JMS and KB edited the paper. FN and KB trained DRGP and WC in use of *in vitro* feeding devices. All authors read and approved the final manuscript.

## Supplementary Figures and Tables

**Additional file 1: Table S1**.

**Ago sequence accession numbers used for phylogenetic reconstruction**.

**Additional file 2: Fig S1.**
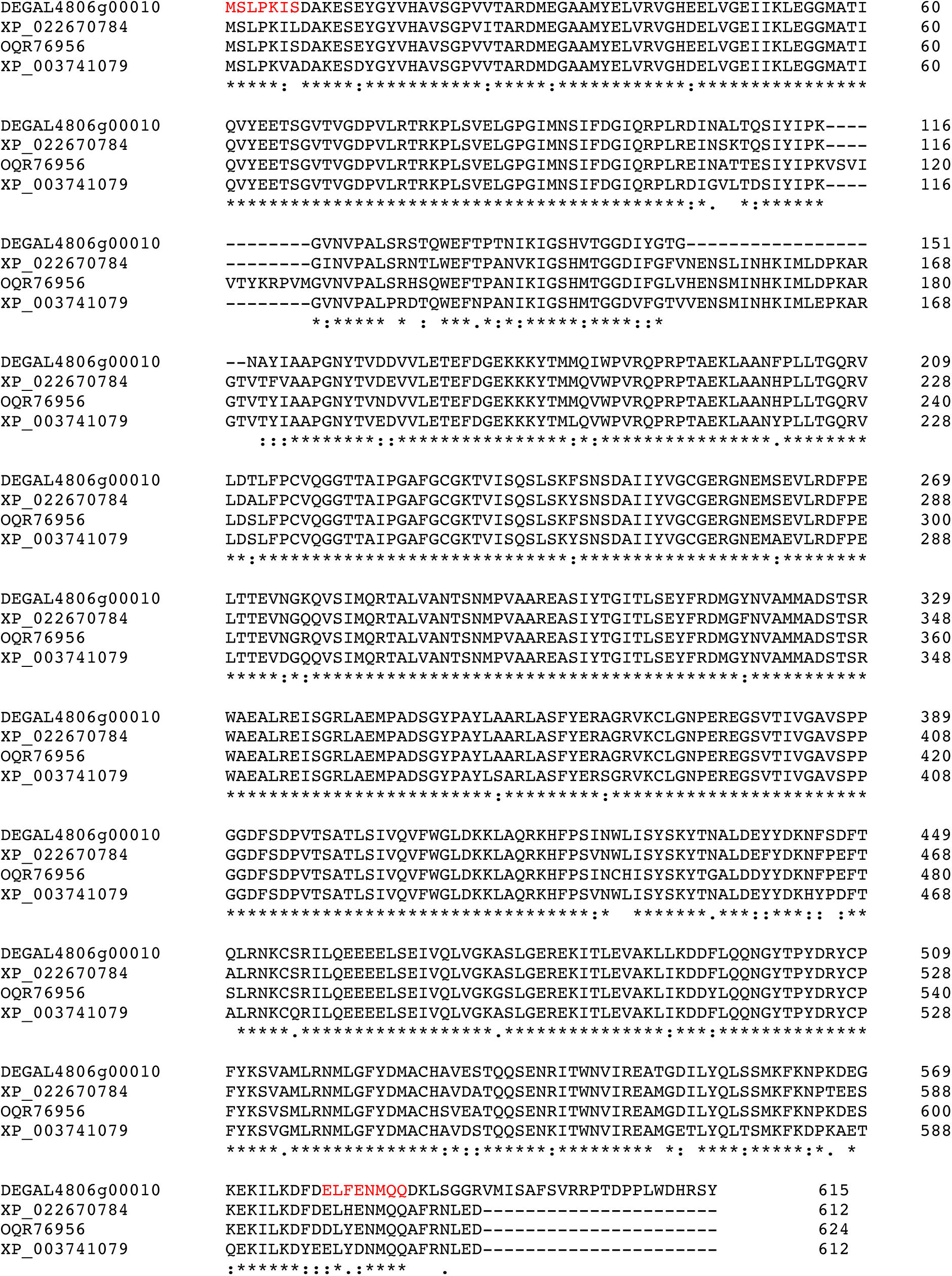
Alignment of mite vATPase A proteins. Dg vATPase A and closely related vATPase A from mites were aligned using MUSCLE. Conserved regions in all sequences are highlighted (*). Sequences included in the alignment are: Dg vATPase A (DEGAL4806g00010); and vATPase A from the following mites: *Varroa destructor* (XP_022670784); *Tropilaelaps mercedesae* (OQR76956) and *Galendromus occidentalis* (XP_003741079). Conserved regions, against which coding sequence amplification primers were designed are highlighted in red.

**Additional file 3: Table S2. qPCR and RNAi construct primer sequences**

**Additional file 4: Fig S2.**
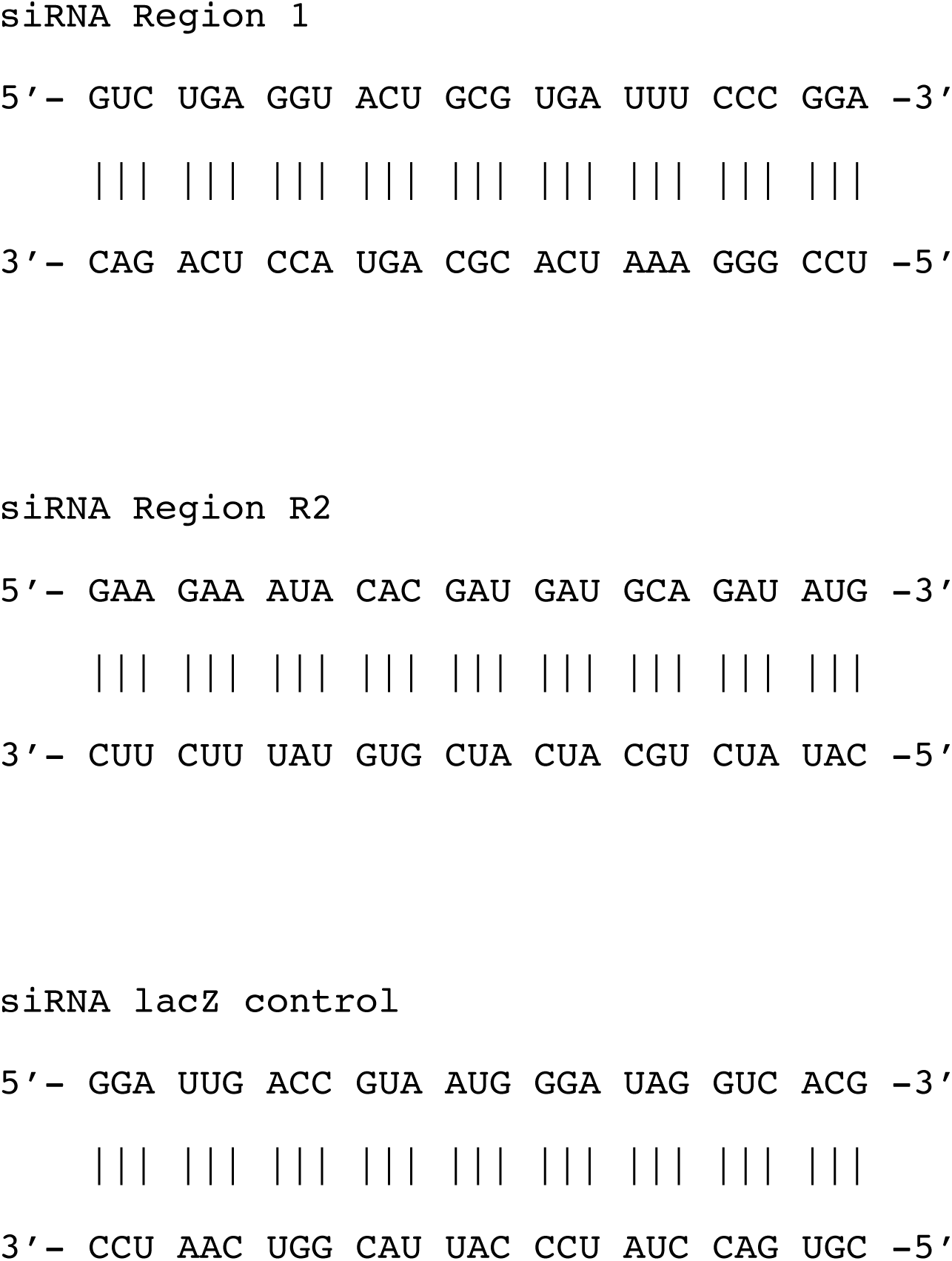
Regions used for synthetic siRNA synthesis.

**Additional file 5: Fig S3.**
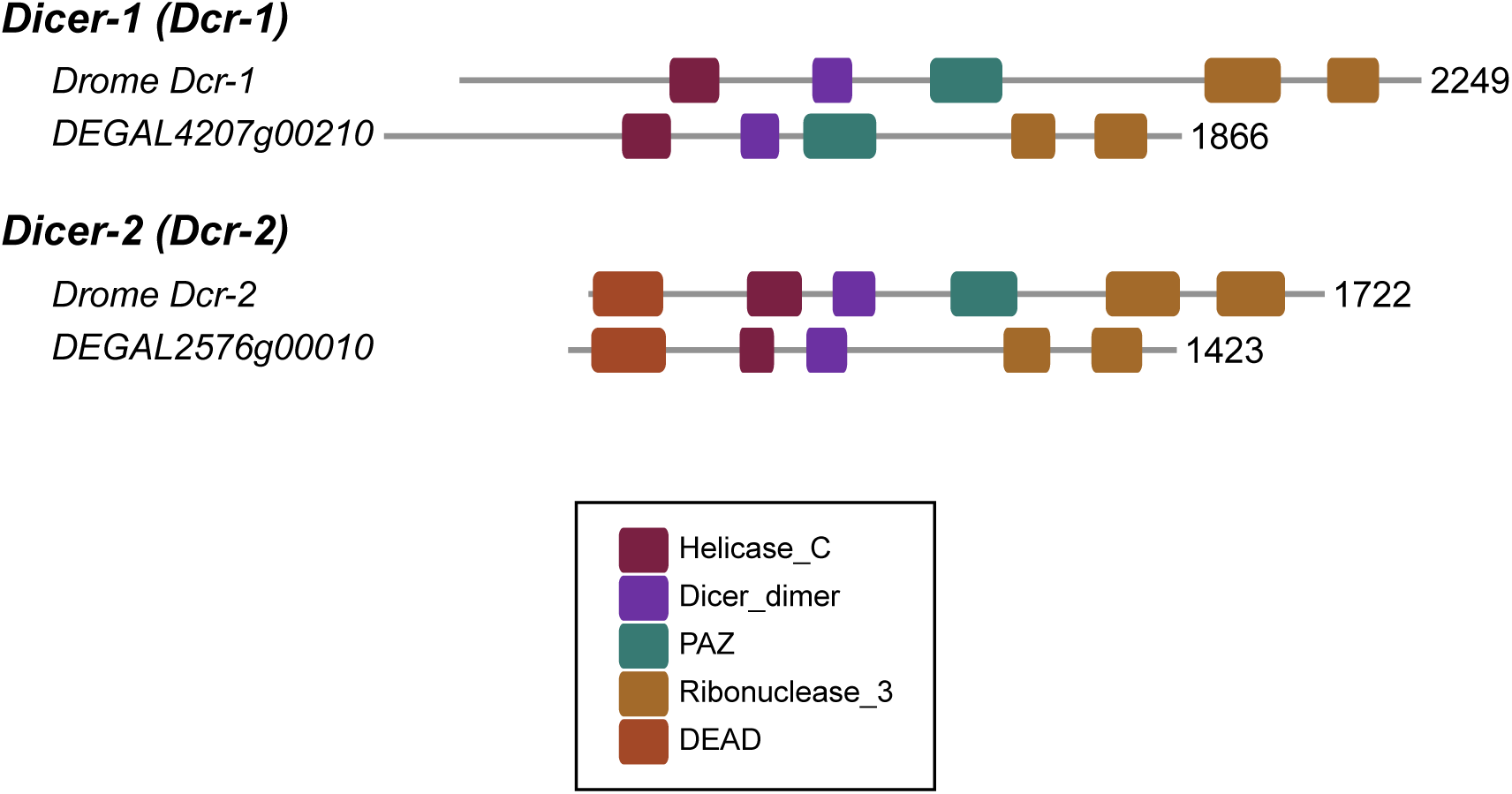
Domain architecture of *D. gallinae* Dicer proteins. For comparison the domain architecture of *D. melanogaster* dicer-1 (Dcr-1) and dicer-2 (Dcr-2) shown. *D. gallinae* Dcr-1 and Dcr-2 were identified as orthologues of Drome Dcr-1 and Dcr-2, respectively. Pfam (https://pfam.xfam.org) functional domains include: Helicase_C (Helicase conserved C-terminal domain, PF00271); Dicer_dimer (Dicer dimerisation domain, PF03368); PAZ (PAZ domain, PF02170); Ribonuclease_3 (Ribonuclease III domain, PF00636); DEAD (DEAD/DEAH box helicase, PF00270). The length of each protein is shown as number of amino acids.

**Additional file 6: Fig S4.**
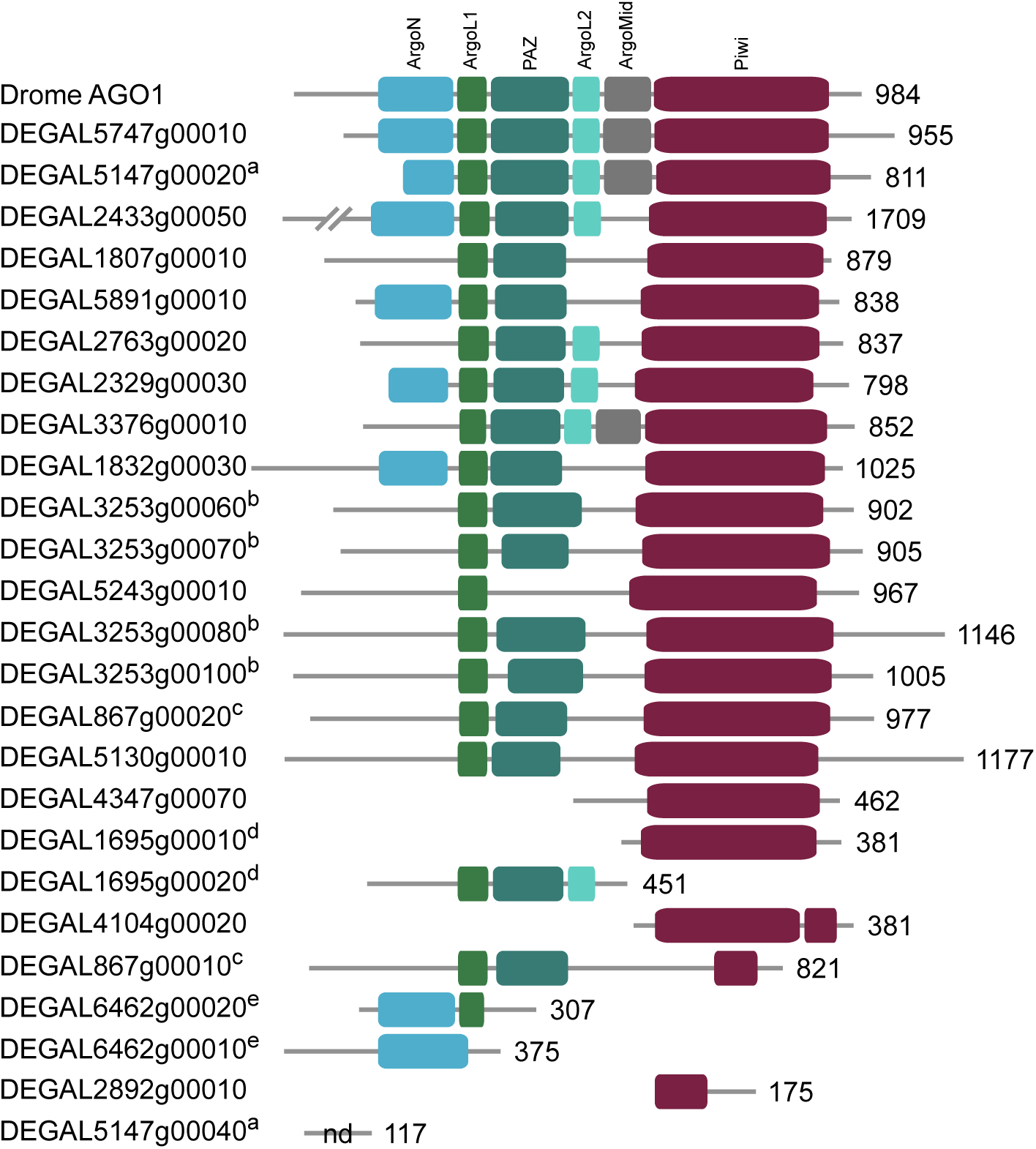
Domain architecture of *D. gallinae* argonaute proteins. For comparison the domain architecture of *D. melanogaster* argonaute-1 (Drome Ago1) is shown. *D. gallinae* argonaute proteins were identified as orthologues of Drome Ago1 and are ranked in order of best blast hit to Drome AGO1. Pfam (https://pfam.xfam.org) functional domains include: ArgoN (N-terminal domain of argonaute, PF16486); ArgoL1 (argonaute linker 1 domain, PF08699); PAZ (PAZ domain, PF02170); ArgoL2 (argonaute linker 2 domain, PF16488); ArgoMid (mid domain of argonaute, PF16487); Piwi (Piwi domain, PF02171). Genes located on the same *D. gallinae* scaffold are highlighted using the superscript letters a-e. The length of each protein is shown as number of amino acids. No Pfam domains were detected (nd, none detected) for DEGAL5147g00040.

